# The transcription factor AtHB23 modulates starch turnover for root development and plant survival under salinity

**DOI:** 10.1101/2021.11.17.468956

**Authors:** María Florencia Perotti, Agustín Lucas Arce, Federico Damián Ariel, Carlos María Figueroa, Raquel Lía Chan

**Affiliations:** Instituto de Agrobiotecnología del Litoral, CONICET, Universidad Nacional del Litoral, FBCB, Colectora Ruta Nacional 168 km 0, 3000, Santa Fe, Argentina

**Keywords:** AtHB23, homeodomain-leucine zipper, HD-Zip I, LAX3, root development, root amyloplasts, salinity stress, starch turnover

## Abstract

AtHB23 is a homeodomain-leucine zipper I transcription factor, previously characterized as a modulator of lateral root initiation and higher-order roots development. The role of this gene in response to salinity stress was completely unknown. To elucidate the role of AtHB23 in response to salinity stress, we combined histochemical β-glucuronidase (GUS) analysis, root phenotyping, starch staining, optic and electronic transmission microscopy, expression studies by RT-qPCR, and transcriptome analysis of silenced, overexpressor, and crossed plants. We revealed that the expression pattern of *AtHB23* is regulated by NaCl in the main and lateral roots, affecting the root phenotype. A severe reduction in primary root length, a significant increment in the initiation of lateral roots, and a low survival rate in salinity conditions were observed in *AtHB23*-silenced plants, whereas *AtHB23* overexpressors showed the opposite phenotype. These developmental defects were explained by the degradation of starch granules and an alteration in starch metabolism. The AtHB23-target gene *LAX3* is repressed in the tip of the main root and affected by NaCl.

We conclude that AtHB23 is vital for plant survival and adaptation to salt stress conditions, and its function is related to the gravitropic response mediated by starch granule turnover, involving the auxin carrier LAX3.

**Highlight:** The transcription factor AtHB23 is crucial for plant survival and adaptation to salt stress conditions, and its function is related to the gravitropic response mediated by starch-granule turnover, involving LAX3.

## Introduction

As sessile organisms, plants must adapt to the soil, climate, and light conditions, as well as to other biotic and abiotic environmental factors in each particular location. Among stressor factors, salinity is one of the major causes of poor crop yields. The situation is further aggravated by undeveloped irrigation strategies, rising population, and pollution from industrial activity. However, plants evolved the capacity to adapt to the soil conditions, displaying various physiological and molecular strategies ultimately integrating external and endogenous signals. Roots are the anchorage organs able to alter their development, extending through the soil to optimize water and nutrient uptake (de Dorlodot *et al*., 2007). Their high plasticity, governed by physiological, genetic, and epigenetic programs, enables the adaptation to changing environments. The integration of various cues allows them to balance growth, development, and stress responses (Schachtman and Goodger, 2008). In this interplay of external and internal signals, different biomolecules, such as hormones and transcription factors (TFs), play crucial roles.

Although phytohormones have been classified as stress or growth-promotion related (Verma *et al*., 2016), numerous studies showed that hormones may participate in alternative pathways and responses, depending on the developmental stage, or environmental condition (Yu *et al*., 2020). Abscisic acid (ABA) is considered the main abiotic stress hormone and has been shown as a crucial player in the responses to drought, salinity, and extreme temperatures, among other factors. Studies about the ABA signaling cascade indicated that this hormone promotes stomata closure which has a high impact on drought response and photosynthesis rate (Vishwakarma *et al*., 2017).

Under salinity stress, in roots, ABA participates in an intricate crosstalk, notably involving auxin. It was suggested that the inhibition of root growth by NaCl is associated with a reduction in auxin content in the tip, mainly due to the alteration of the auxin carriers AUX1 and PIN2 (Liu *et al*., 2015). The development of lateral roots (LR) is also affected by salinity, exhibiting an enhanced growth under low NaCl concentration (≤50 mM) and repressed at higher concentrations. Notably, these effects are dependent on auxin transport and distribution (Zolla *et al*., 2010).

In early root developmental stages, auxin is transported toward the tip through the central cylinder (Friml *et al*., 2003) by specific carriers (AUX1, LAX1, LAX2, and LAX3), which exhibit differential expression patterns (Péret *et al*., 2012; Swarup and Péret, 2012). Among these carriers, AUX1 has been linked to LR initiation and LAX3 to LR emergence (Marchant *et al*., 1999, 2002). Efflux/influx auxin carriers are also involved in root gravitropism, which depends on a differential hormone gradient between the upper and lower sides of the root (Zhang *et al*., 2019). This gradient regulates the synthesis of starch granules, a process severely affected by salinity (Korver *et al*., 2020; Zhang *et al*., 2019). In this way, the gravitational potential energy is converted into a biochemical signal involving statoliths, which are compact starch aggregates formed in the columella cells (Leitz *et al*., 2009). Besides the gravitropic response, roots also display hydrotropism, interfering with gravitropism in a complex and unknown way. During the hydrotropic response, amyloplasts are degraded in the columella cells reducing the gravitropic response (Nakayama *et al*., 2012).

The complex molecular events modulating root architecture and plasticity involve TFs playing different roles in the adaptation to the environment. TFs families involved in these processes include ARF (Auxin Response Factors), LBD (Lateral Organ Boundaries (LOB) Domain), WRKY (named after the four common WRKY amino acids and a zinc finger-like motif), MYB (MYeloBlastosis oncogene), HD (Homeodomain), bHLH (Basic Helix–Loop–Helix), NAC (NAM, ATAF, and CUC), AP2/ERF Domain (APetala 2/Ethylene-Responsive element-binding Factor), MADS (named after MCM1; AGAMOUS; DEFICIENS, and SRF members), GRF (Growth-Regulating Factor), and GRAS (named after the first three members GIBBERELLIC-ACID INSENSITIVE (GAI), REPRESSOR of GAI (RGA) and SCARECROW (SCR) members (González, 2016; Hong, 2016), which have been functionally characterized in various plant species (T Li *et al*., 2020; Renau-Morata *et al*., 2020). Notably, members from these families are interconnected through protein-protein or protein-DNA interactions to activate full transcriptional programs. In particular, members of the ARF and LBD families have been identified as master regulators of LR development, and several among them also mediate the crosstalk between hormones (Friml *et al*., 2003; Xu *et al*., 2020; Banda *et al*., 2020; Lavenus *et al*., 2013; Lee *et al*., 2009).

The homeodomain-leucine zipper family (HD-Zip) is unique to the plant kingdom. The members of this family have been classified into four subfamilies according to structural and functional features (Capella *et al*., 2015). Many of them participate in root development (reviewed by Perotti *et al*., 2021). Particularly, HD-Zip I TFs were associated with abiotic stress responses, although most studies have been focused on the aerial organs of the plant (Perotti *et al*., 2021).

AtHB23 is a transcription factor belonging to the HD-Zip I subfamily, and it was described as a modulator of LR initiation through the direct regulation of *LAX3* and *LBD16*. This gene is regulated by ARF7/19 (Perotti *et al*., 2019, 2020). According to publicly available databases, the expression of *ARF19* is affected by salinity (http://bar.utoronto.ca/; Dinnery *et al*., 2008), and because HD-Zip I TFs were associated with abiotic stress (Perotti *et al*., 2017), we wondered whether AtHB23 participates somehow in the response to salinity in Arabidopsis roots.

In this work, we show that *AtHB23* expression is regulated by NaCl, both in primary and LRs. *AtHB23*-silenced (*amiR23*) plants exhibit an enhanced sensitivity to NaCl treatments affecting the gravitropic response and survival. Accordingly, starch granules were degraded when plants were treated with NaCl and were unable to recover once transferred back to normal conditions. Transcriptomic analyses revealed that AtHB23-dependent reprogramming in response to salt includes the regulation of starch turnover related genes. Moreover, we demonstrate that the ATHB23 target gene *LAX3* is also regulated by salinity and participates in the root adaptive response.

## Materials and methods

### Genetic constructs

*35S::amiR23, 35S::AtH23, prAtHB23:GUS, prLBD16:GUS, prLAX3:GUS*, and *35S::amiR23 x prLAX3:GUS* were previously described (Ribone *et al*., 2015; Perotti *et al*., 2019). *prAtHB23Δ1273:GUS:GFP*: a PCR reaction using specific oligonucleotides (Table S1) and genomic DNA was carried out. The amplicon (1273 bp upstream the ATG) was cloned into the *pENTR3C* plasmid using the *BamH1* and *EcoRI* restriction sites. After that, a Gateway® (Invitrogen) recombination was performed in the destination *pKGWFS7*vector. The resulting construct was called *prAtHB23*_*S*_:*GUS*.

### Plant material, growth conditions, and transformation

For NaCl treatments and root phenotyping, Arabidopsis thaliana plants (Col 0 ecotype) were grown in a growth chamber at 22–24°C under long-day conditions (16/8 h light/dark cycles) with a light intensity of approximately 70 µmol m^-2^sec^-1^ in vertical square Petri dishes (12 × 12 cm) with Murashige–Skoog medium supplemented with vitamins (MS; PhytoTechnology Laboratories, https://phytotechlab.com/home).

### Histochemical GUS staining

Plants were immersed in GUS staining buffer (1 mM 5-bromo-4-chloro-3-indolyl-GlcA in 100 mM sodium phosphate, pH 7, 0.1% (v/v) Triton X-100, and 100 mM potassium ferrocyanide), vacuum was applied for 5 min, and then plants were incubated at 37°C for 12 h. Chlorophyll was cleared from the plant tissues by immersion in 70% ethanol.

### Root phenotyping

Seeds were surface sterilized and placed 1 cm from the top for 3 days at 4°C before placing the dishes in the growth chamber. The lengths of the main root were measured using the RootNav free software from photographs of the plates (Pound *et al*., 2013).

### Salinity treatments

Assays done with *prAtHB23:GUS* plants: seedlings were grown in normal conditions in vertical Petri dishes (12 × 12 cm) an then placed in plates with the same medium supplemented with NaCl (concentrations indicated in each Figure legend). LR and LRP observation and counting were done with seedlings grown in normal conditions during 3 days and placed in plates supplemented with NaCl for additional 5 days.

The survival experiment was performed with plants placed in 100 mM NaCl after three days of growing in normal conditions.

To amyloplast observation and transcript quantification, 5-day-old plants grown in control conditions were treated during 8 h with 150 mM NaCl. After that, half of the plants were transferred to control conditions and the other half remained in the salinity medium during 72 additional hours.

### Amyloplasts Staining and Light Microscopy Observation

To observe the amyloplasts in the columella cells of the root tips, 15-20 Arabidopsis roots (5 day-old) were dipped in Lugol staining solution (Sigma-Aldrich) for 8-10 min, and then observed in an Eclipse E200 Microscope (Nikon, Tokyo, Japan, https://www.nikon.com/) equipped with a Nikon Coolpix L810 camera.

### RNA isolation and analysis

The total RNA used in quantitative reverse transcriptase RT-qPCR was isolated from Arabidopsis roots using Trizol reagent (Invitrogen, https://www.thermofisher.com/ar/), according to the manufacturer’s instructions. One microgram of RNA was reverse transcribed using oligo(dT)18 and M-MLV reverse transcriptase II (Promega, https://worldwide.promega.com/). Quantitative real time PCR (qPCR) was performed using a StepOnePlus Real-Time Systems of Applied Biosystems TM. Each reaction contained 10 μl final volume having 5 μl Taq TM SYBR^®^ Green Supermix, 0,2 μl of each specific oligonucleotide (10 pmol/μl; Table S1) and 1/20 of RT product. Fluorescence was continuously recorded at 72°C during 40 cycles. The quantification of mRNA levels was achieved by normalization against ACTIN transcripts levels (*ACTIN2* and *ACTIN8*), following the *ΔΔ*Ct method. All reactions were performed with at least three biological replicates and bars represent SEM.

### Histology and electronic microscopy

The roots were fixed in glutaraldehyde at 2% (v / v) in 0.2 mM potassium phosphate buffer (pH 7.4) and kept at room temperature for 2 h (under vacuum) and then brought to 4°C overnight. Segment were rinsed with the same buffer and subsequently fixed in 2% (w / v) OsO_4_ in buffer (pH 7.0) at room temperature for 2 h (under vacuum), and then dehydrated in a series of acetone and embedded in Durcupan resin (whit vacuum application at each step). For light microscopy, sections about 0.5 µm thick were stained with 0.05% (w / v) toluidine blue in bidistilled water. For transmission electron microscopy, ultrathin sections (70-90 nm in thickness) were stained with uranyl acetate and lead citrate and observed under a Philips EM201 transmission electron microscope (TEM) at an acceleration voltage of 60-80 kV.

### Transcriptome analysis

For transcriptome analyses, Col 0 and *amiR23* seeds were grown for 3 days in normal conditions. After that one half of the plates were transferred to new MS plates supplemented with 75 mM NaCl and samples were harvested 6 days after that. Total RNA was isolated from roots. The experiment was performed with biological replicates of each genotype/treatment. Isolated RNA was lyophilized before shipping.

Sequencing was performed by the BGI Genomics Sequencing Service (https://www.bgi.com/global/), using the DNBseq platform, and obtaining 100 nucleotide long paired-end reads. Raw reads were first quality trimmed and filtered with Trimmomatic (version 0.36; Bolger *et al*., 2014) and then aligned to the *Arabidopsis thaliana* genome (TAIR10) using STAR (version 2.5.2b, Dobin *et al*., 2014), which was guided by the gene and exon annotation from Araport (V11 201606, Pasha et al., 2020). Samtools (version 1.8; Li *et al*., 2009) was then used to keep only primary alignments with a minimum MAPQ of 3. Read quality before and after trimming was analyzed with FastQC (version 0.11.5; Andrews 2010) and, together with mapping efficiency, it was summarized with MultiQC (version 1.7; Ewels *et al*., 2016). Read counts on each gene were calculated with featureCounts (version 1.6.2; Liao *et al*., 2014). This pipeline was run with the aid of the Snakemake workflow engine (Köster and Rahmann, 2012). Differentially expressed genes were determined with DESeq2 (Love et al., 2014; R Core Team, 2020) filtering out genes with counts below 10 in all samples, and combining factors (i.e. genotype, WT or *amiR23* and treatment, control or salinity) into a single factor that was used in a simple design formula (as recommended by the package vignette as alternative to more complex models).

Sequencing data were deposited at the European Nucleotide Archive (https://www.ebi.ac.uk/ena/) with the accession id PRJEB47043.

### Statistical analyses

The data shown in 1, 2, 3, and S5 were analyzed using a two-way ANOVA considering genotype and treatment. When interaction terms were significant (P < 0.01), differences between means were analyzed using Tukey comparison and are indicated by different letters. The number of biological replicates for each assessment are indicated in the corresponding figures.

The Venn diagram in Fig. 4A was performed using the R package VennDiagram (Chen, 2018) and the heatmap in Fig. 4B was done with the pheatmap package (Kolde, 2019).

### Accession numbers

AT1G26960 (AtHB23); AT2G42430 (LBD16); AT1G77690 (LAX3)

## Results

### *AtHB23* expression is regulated by NaCl strongly affecting root architecture

AtHB23 has been identified as a crucial player in LR initiation. This gene is a direct target of ARF7/19, regulated by salinity (Perotti *et al*., 2019; http://bar.utoronto.ca/; Dinnery *et al*., 2008). Since primary root and LR development are closely linked to soil conditions, we tested whether the role of AtHB23 was affected by salinity using NaCl. Root phenotype was evaluated in *AtHB23* overexpressors (AT23), knock-down (*amiR23*), and wild type (Col 0) 3-day-old plants treated for additional 5 days with different NaCl concentrations. Both silenced and overexpressors showed an altered phenotype compared to controls. The differences were more pronounced concomitantly with growing concentrations of NaCl. Knock-down plants exhibited a significantly reduced primary root length in 50 and 75 mM NaCl (Fig. 1A), whereas AT23 overexpressors showed the opposite phenotype in control conditions and 50 mM NaCl (Fig. 1E). When the concentration reached 75 mM, the difference between the WT and AT23 was undetectable (Fig. 1E). LRP density was higher in *amiR23* seedlings, in agreement with the previously reported results (Perotti *et al*., 2019), but this difference with the WT diminished at increasing concentrations of NaCl (Fig. 1B). In contrast to control conditions, AT23 plants exhibited lower LRP density than in the WT in response to NaCl (Fig. 1F). LR density was larger in AT23 plants than in controls in 75 mM NaCl, whereas silenced plants, at this NaCl concentration, behaved similarly to AT23 (Figs. 1C and 1G). Total LR followed the pattern of lateral root primordium (LRP), suggesting more influence on the LRP number than on LRs (Figs. 1D and 1H).

**Figure 1.**
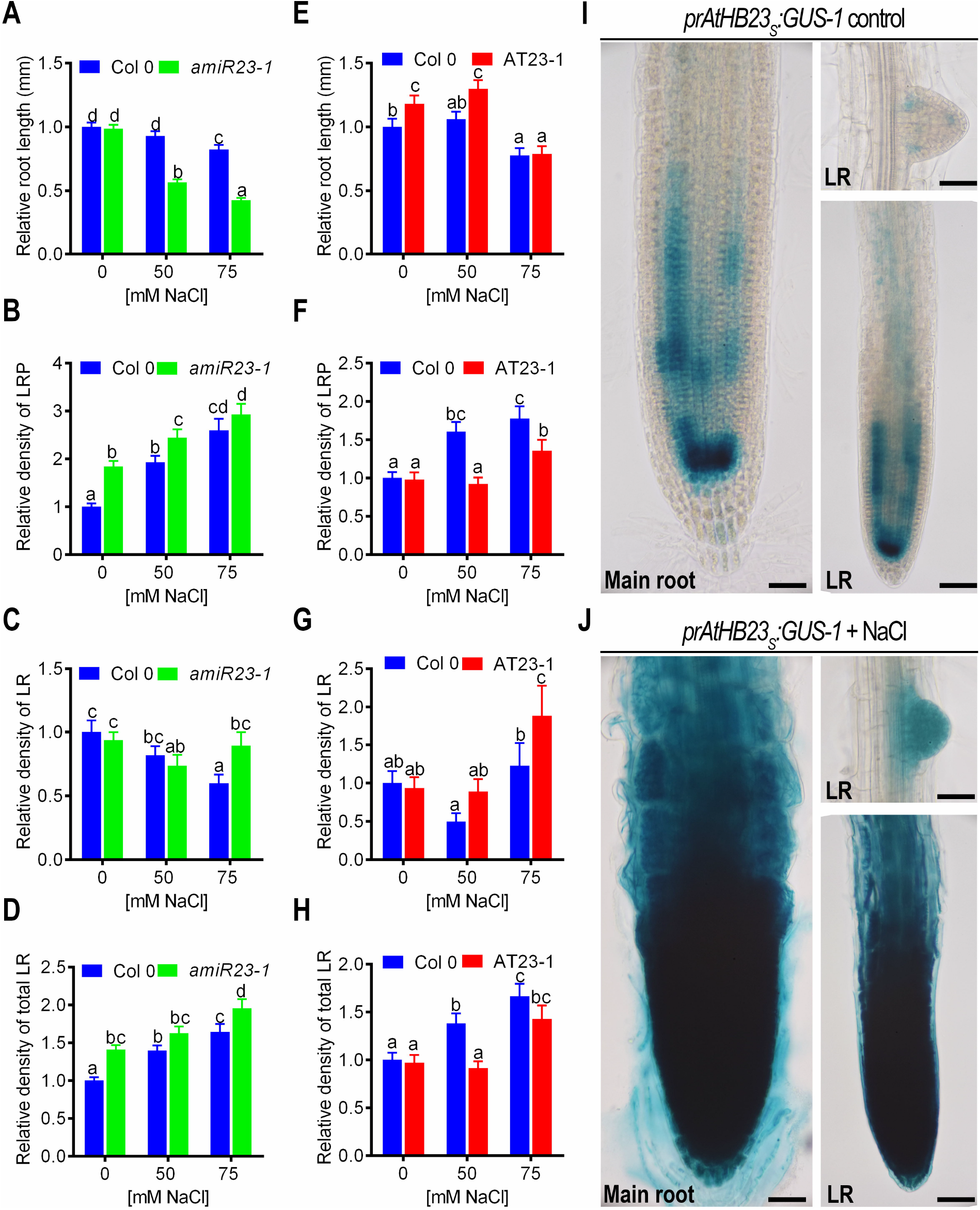
The expression pattern of *AtHB23* is differentially affected by NaCl in the main and lateral roots, altering whole root development. (**A-D**) Relative primary root length (**A**) of8-day-old Col 0 and *amiR23-1* silenced plants, relative density of lateral root primordium (LRP, **B**), lateral root (LR, **C**), and total LR (LRP + LR, **D**) calculated as the number of LRP or LR/mm of main primary root, grown in control conditions during 8 days (control) or 3 days and treated with NaCl (50 or 75 mM), during 5 additional days. (**E-H**) The same as in (**A-D**) evaluated in Col 0 or AT23 *(AtHB23* overexpressor) plants. The values were normalized with those measured in the Col 0 control, taken as 1 (one). All the assays were repeated at least three times with N: 15/genotype. (**I**) Illustrative pictures showing GUS expression driven by *AtHB23* promoter (short version, *prAtHB23*_*s*_*:GUS)* in the tips of main and lateral roots of 8-day-old seedlings grown in control conditions. (**J**) The same as in (**I**) in plants grown 3 days in normal conditions and then placed during 5 additional days in 75 mM NaCl. LR are emerged lateral roots. Black bar represents 50 µm.

To evaluate the expression pattern of *AtHB23* in salinity conditions, transgenic plants carrying the construct *prAtHB23*_*L*_*:GUS* (1793 bp upstream from the start codon) were treated with NaCl and GUS activity visualized by histochemistry. In normal conditions, the promoter activity was restricted to the base of LRP at stages V and VI and disappeared when the LR emerged, whereas in advanced stages of emerged roots, a signal in the vascular system of both the main and LR was observed (Fig. S1A). After treating the roots with NaCl, GUS staining was restricted to the LRP and disappeared or diminished from the vascular system of emerged roots (Fig. S1B).

Given the altered phenotype observed in plants with reduced (*amiR23*) or augmented (AT23) levels of *AtHB23*, we wondered how this TF was affecting the development of the main root when its expression was not detectable in the root tip. Regulatory elements can be found in various locations, near or far away from coding sequences. Considering that the promoter region we isolated was arbitrarily defined as the 1793 bases upstream of the start codon (Perotti *et al*., 2019), we decided to dissect the *AtHB23* promoter region to uncover its potential activity in the root tip. To this end, we cloned shorter segments fused to the reporter gene *GUS* and transformed WT plants. Interestingly, a segment of 1273 bp showed a strong activity in the tips of the main and secondary roots, and in the emerged LR (Figs. 1I and S2). Since this segment was 520 bp shorter than the one previously assessed (Perotti *et al*., 2019, 2020 and Fig. S1), our results suggest that the deleted region might include a repressor box. Remarkably, when these plants were subjected to NaCl treatments, GUS staining was significantly increased in the tip of the main root, in recently emerged LR, as well as in the tip of secondary LR (Figs. 1J and S2). Therefore, the expression pattern of the shorter *AtHB23* promoter in the root tip in response to NaCl was in agreement with the differential root growth observed in *AtHB23*-deregulated plants.

### Long treatments at increasing NaCl concentrations have a severe effect on *amiR23*-silenced plants

To further characterize root architecture in salinity conditions, seedlings were subjected to increasing times and NaCl concentrations. The difference between WT, AT23, and *amiR23*-silenced plants in primary root length slightly increased between 4 and 10 days after sowing (Fig. 2A). However, the difference in the main root growth became more evident when seedlings were treated with NaCl. In 25 mM NaCl, the difference between genotypes augmented significantly (Fig. 2B) whereas, in 50 and 75 mM NaCl, *amiR23* plants almost arrested their development while AT23 and WT seedlings maintained a similar growth rate with a slight advantage for AT23 (Figs. 2C and 2D). Notably, the primary roots of *amiR23* seedlings showed a curved shape, suggesting that the gravitropic response was affected (Fig. 2).

**Figure 2.**
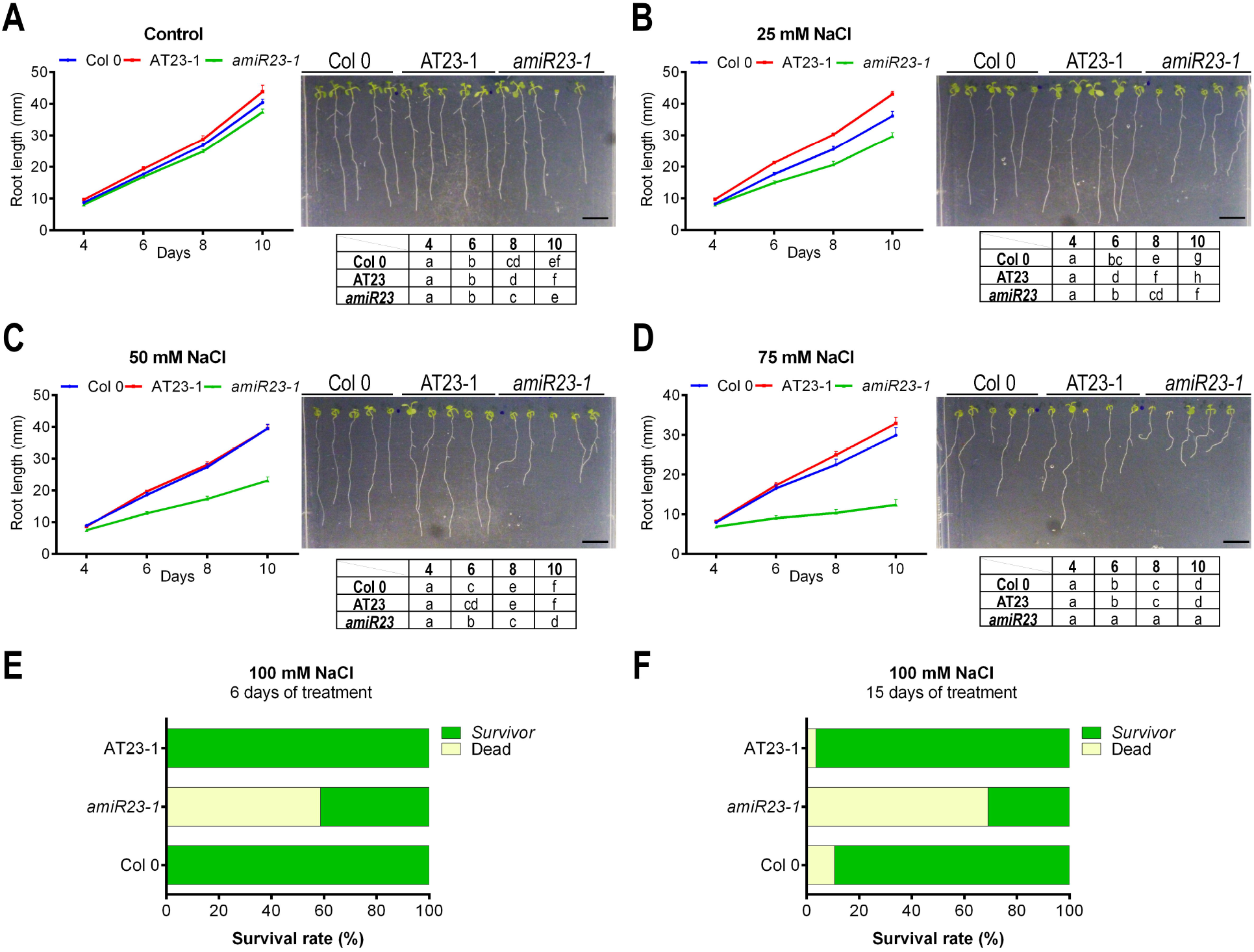
AtHB23 promoted main root elongation in front of salt stress. (**A**) Main root length evaluated in Col, AT23-1 (overexpressors), and *amiR23-1* (silenced) seedlings grown in normal conditions. In the left panel quantitative measurements performed from day 4 after sowing and until day 10, and in the right panel, illustrative pictures of the same roots taken the 10th day. The same plants as in (**A**) were transferred 3 days after sowing to plates supplemented with 25 mM (**B**), 50 mM (**C**) or 75 mM (**D**) NaCl. The assays were repeated at least three times with N: 15/genotype. Black bar represents 1 cm. Different letters indicate significant differences (Tukey test, P < 0.01). Error bars represent SEM. (**E-F**) Survival rate of plants (Col 0, AT23-1 and *amiR23-1)* placed in plates with 100 mM NaCl, 3 days after sawing during the 6 (**E**), and 15 (**F**) additional days. Yellow columns indicate the % of dead plants whereas green ones the % of survivors. The assays were repeated at least three times with N: 15/genotype.

Strikingly, the assessment of the survival rate of the different *AtHB23* knock-down and overexpressor lines at 100 mM NaCl for 6 and 15 days revealed a boosted sensitivity of *amiR23* plants to high salinity. After 6 days of treatment, 60 % of *amiR23* plants died, whereas all WT and AT23 plants survived. After 15 days, a growing percentage of WT plants died whereas most of AT23 plants still survived, in contrast to the low survival rate of *amiR23* plants (Figs. 2E and 2F).

### The *amiR23*-silenced plants lost the starch granules in the root tip after salinity treatment

To better characterize the high sensitivity of *amiR23* plants to NaCl, we grew them together with the WT and AT23 genotypes during five days in normal conditions. Then, seedlings were transferred to the same medium containing 150 mM NaCl for eight additional hours. Afterwards, a group of plants was transferred back to MS medium to recover, while another group remained in the NaCl-containing medium. From those plants kept in NaCl, notably WT and AT23 slowly adapted and recovered similarly to those transferred to the NaCl-free MS medium. On the other hand, the *amiR23* plants kept in NaCl died. The roots of plants from all groups were treated with Lugol staining solution to visualize starch in their root tips. In normal conditions, AT23 plants exhibited more starch than the WT, whereas the *amiR23*-silenced plants had significantly less starch than the controls (Figs. 3A and 3C). After NaCl treatment, starch accumulation was affected in all the genotypes but AT23 plants showed a better performance than control plants, whereas it was not possible to visualize the stained polyglucan in *amiR23* tips (Figs. 3A and 3C). Notably, after recovery in a normal medium, both WT and AT23 roots exhibited restored amyloplasts, whereas *amiR23*-silenced plants did not recover and continued to lack visible starch (Fig. 3A). The same experiment was carried out with independent lines of *amiR23*-silenced plants showing similar characteristics to those of *amiR23-1*. Moreover, the phenotype of *amiR23*-silenced plants was rescued by crossing them with AT23 plants (*amiR23* × AT23 genotype), restoring the ability to recover from NaCl treatment (Fig. S4).

**Figure 3.**
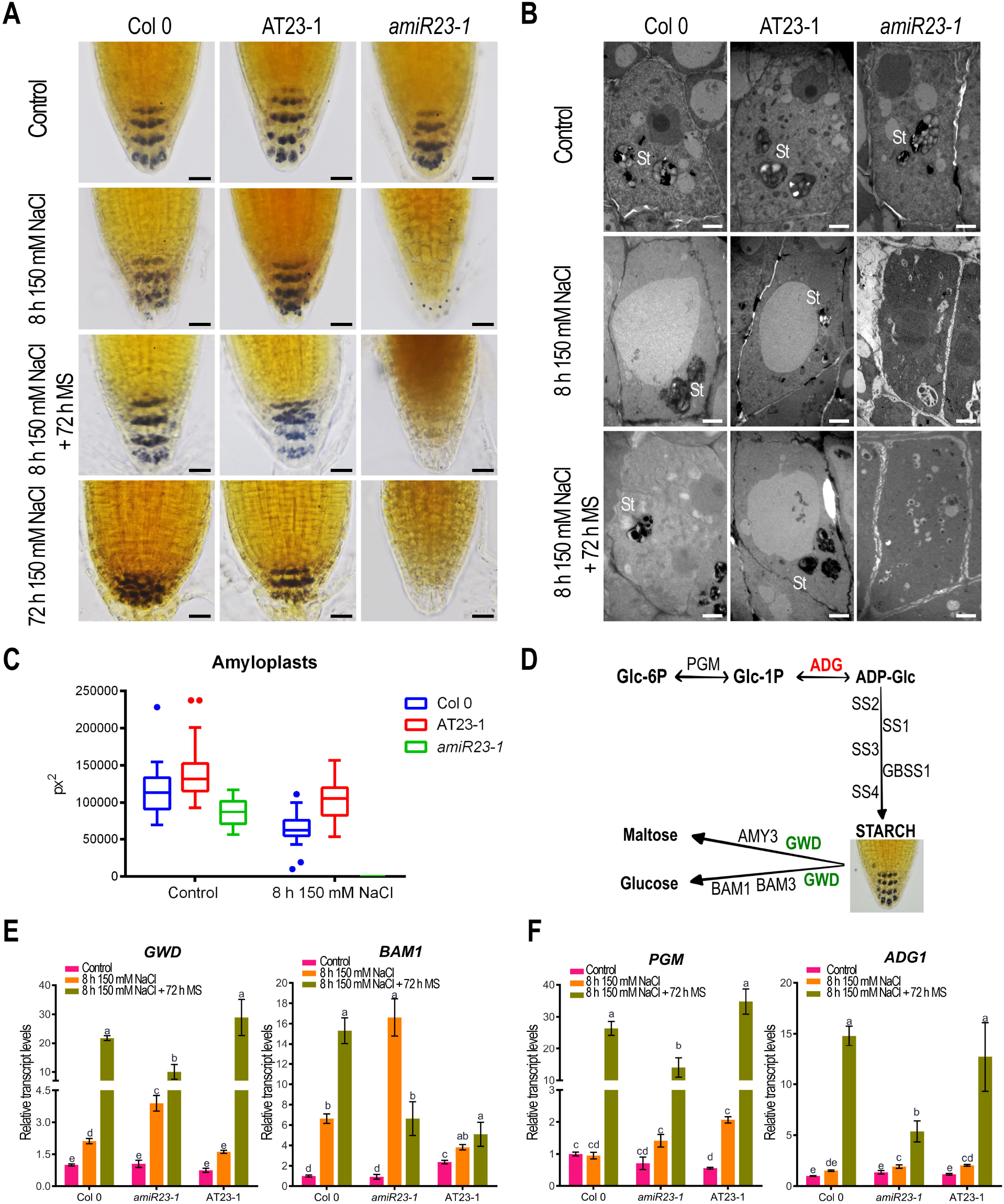
The amyloplasts in the root tip of *amiR23* silenced plants were degraded in NaCl medium and were unable to recover. (**A**) From top to bottom, illustrative pictures of root tips (5-day-old) stained with Lugol solution of Col 0, AT23, and *amiR23* seedlings grown in normal conditions (top), after 8 (upper-middle) or 72 h (bottom) treatment with 150 mM NaCl, and then transferred to normal conditions for additional 72 h (lower-middle). Black bar represents 50 µm. (**B**) Illustrative pictures taken with Transmission electron microscopy (TEM) of root tips of 8-day-old seedlings grown in normal conditions (top), treated during 8 h with NaCl (middle), and placed again in normal conditions for 72 h additional hours (bottom). White bars represent 2 µm. (**C**) Starch quantification by ImageJ software in the roots treated or not with 150 mM NaCl. (**D**) Schematic representation of the metabolic pathways for starch synthesis or degradation. (**E-F**) Transcript levels of key genes in Col 0, AT23-1, and *amiR23-1* plants grown in normal conditions during 5 days, treated 8 h with150 mM NaCl and placed to recover in MS medium during additional 72 h. Genes assessed participating from degradation were *GWD* and *BAM1* (**E**), and from synthesis *PGM*, and *ADG1* (**F**). All the values were normalized with the one obtained in Col 0. Bars represent SEM. Data were analyzed using a two-way ANOVA considering genotype and treatment. Different letters indicate significant differences (Tukey test, P < 0.01). The accession numbers of the tested genes are the following: AT5G48300 (ADG1), AT5G51820 (PGM), AT1G10760 (GWD), AT3G23920 (BAM1).

To elucidate whether *amiR23* plants lost starch or if the amyloplasts were degraded, resulting in starch dispersion, then not visible under the optic microscope, a transmission electronic microscopy (TEM) was used to observe the treated roots. TEM images indicated that *amiR23* plants neither had amyloplasts nor disseminated starch after the NaCl treatment (Fig. 3B). Moreover, amyloplasts did not reappear after transferring back the plants to a normal medium (Fig. 3B).

Altogether, our results indicate that AtHB23 is a key factor participating in the adaptation of Arabidopsis plants to high salinity, notably assuring starch turnover and/or amyloplast biogenesis.

### Starch synthesis and degradation are significantly affected by AtHB23

We then wondered if starch biosynthesis, its degradation, or both were affected upon *AtHB23* deregulation. To this end, we evaluated key genes involved in the pathways of starch biosynthesis and degradation in AT23 and *amiR23* silenced plants (Fig. 3D). Collectively, our results showed an accumulation of transcripts encoding enzymes participating in starch degradation and reduced levels of transcripts encoding enzymes necessary for the synthesis of the polyglucan (Figs. 3E and F). For example, *GWD* and *BAM1* encoding the enzymes degrading starch to maltose and glucose (Fig. 3D), were strongly induced in *amiR23* roots subjected to NaCl for 8 h (Fig. 3E). In agreement with this, *PGM* and *ADG1* involved in starch biosynthesis (Fig. 3D), were induced in all genotypes after recovery, but significantly more in WT and AT23 plants than in the *amiR23* silenced plants (Fig. 3F). Other genes, also involved in starch synthesis and degradation, showed significant changes in their transcripts depending on *AtHB23* levels. For example, *SS2, SS3, SS4*, and *GBSS1*, which participate in the pathway of starch synthesis (Fig. 3D), were significantly induced after recovery from NaCl treatment in MS medium but such increase was lower in the silenced plants (Fig. S5). Differently, *SS1* was strongly induced in the *amiR23* silenced plants after NaCl treatment and showed an upregulation tendency in the WT, whereas in the AT23 genotype the differences were not significant (Fig. S5). Finally, *AMY3* and *BAM3*, involved in starch degradation (Fig. 3D), were upregulated after recovery in all the genotypes but to a lesser extent in the silenced plants (Fig. S5). Altogether, these results indicated that starch turnover was significantly altered by the levels of *AtHB23*. However, this fact solely cannot explain the absolute lack of starch observed in *amiR23* plants, indicating that other mechanisms must be modulating this process.

### A transcriptome analysis strongly supported the role of AtHB23 in carbohydrate turnover

Because of the crucial role played by AtHB23 in salinity conditions, we wondered which other signal transduction pathways, aside from starch turnover, could be modulated by this TF. To further investigate this, an RNA-Seq transcriptomic experiment was carried out using 9-day-old WT and *amiR23*-silenced plants grown in control conditions or treated with 75 mM NaCl. In contrast to the analysis performed for a subset of genes in response to short treatments with NaCl, this transcriptomic approach served to uncover the AtHB23-dependent transcriptional reprogramming during the plant adaptation to salinity. In control conditions, only 24 genes significantly changed their expression in *amiR23* versus WT plants (Fig. 4A). Interestingly, the number of differentially expressed genes (DEGs) between genotypes increased to 2025 in salinity conditions (Fig. 4A), comparing to control conditions, the salinity treatment resulted in 2775 DEGs in WT plants and 4225 DEGs in *amiR23* plants. Thus, AtHB23 appears as a central regulator of the adaptation to long-term stress.

**Figure 4.**
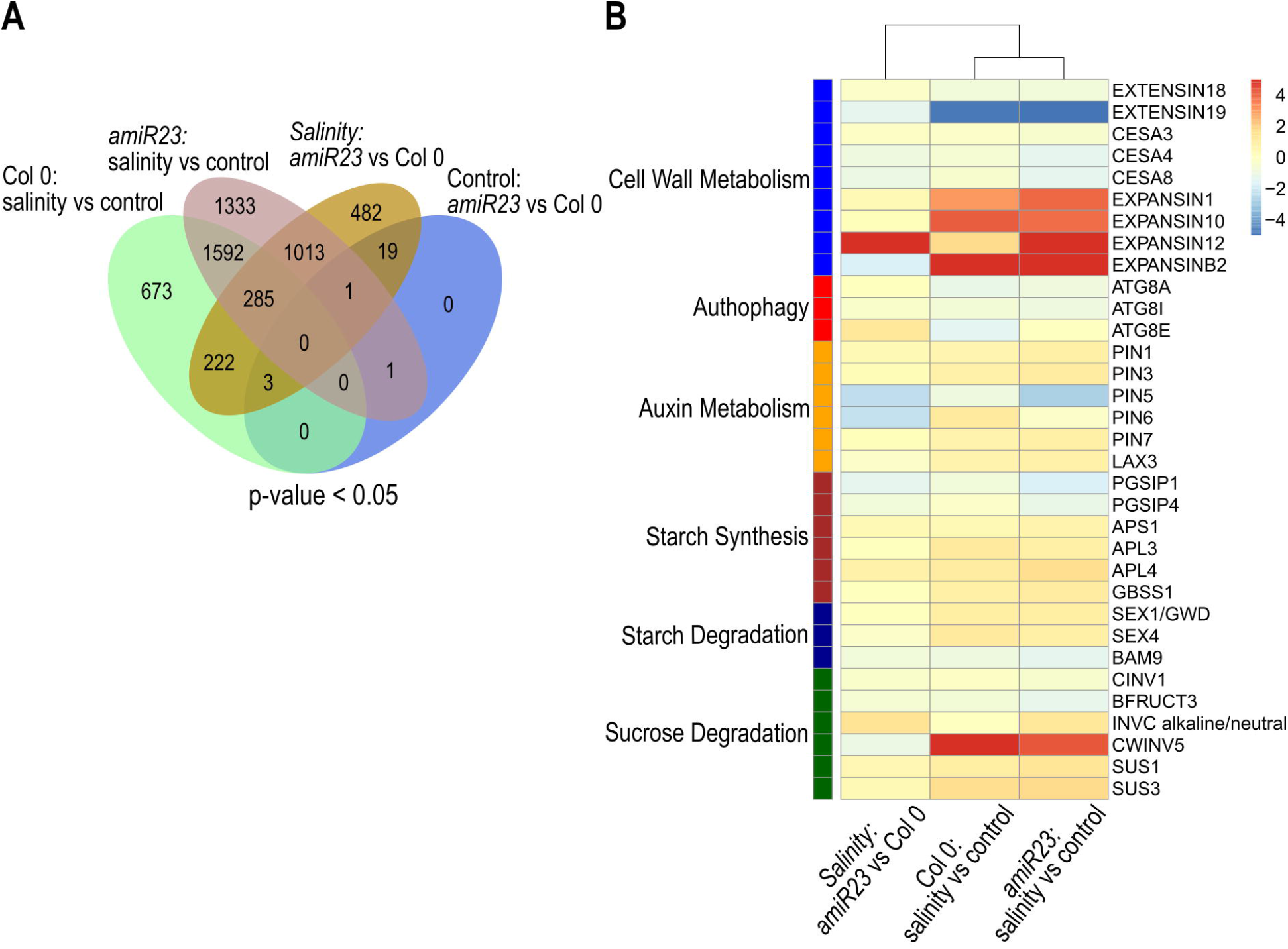
Transcriptome analyses indicated a differential regulation in *amiR23* roots of genes involved and starch metabolism in front of salinity. (**A**) Venn Diagram showing the overlap of the sets of differentially expressed genes (FDR-adjusted p-value < 0.05) in the four different contrasts analyzed, in which the level of one factor is fixed and the two levels of the other factor are compared, e.g. the genotype is WT and the comparison is performed between salinity and control conditions. (**B**) Heatmap showing the expression pattern of relevant genes (rows) in relation to the *amiR23* phenotype. They were selected based on the results of the Mapman analysis of DEGs (Fig. S6), and the inspection of the complete transcriptome (Table S2). The functional annotation of genes is highlighted on the left. Only the three contrasts with a relevant number of DEGs were included (columns).

An analysis performed with the Mapman software allowed the identification of the pathways in which DEGs were involved.

We first compared results obtained for *amiR23*-silenced plants treated with NaCl versus control conditions. There were major changes in transcripts related to lipids, secondary metabolism and cell wall (Figs. 4B and S6A). Some of these were severely reduced, such as *EXTENSIN18* and *EXTENSIN19*, which encode hydroxyproline-rich proteins putatively involved in cell wall assembly. Also, several transcripts encoding different cellulose synthase isoforms (*CESA3, CESA4*, and *CESA8*) were decreased (Fig. 4B and Table S2). Conversely, some transcripts encoding extensins (which mediate cell wall loosening) were highly increased, including EXPANSIN1, *EXPANSIN10, EXPANSIN12*, and *EXPANSINB2* (Fig. 4B and Table S2). Intriguingly, there was a massive increase of transcripts encoding components of the light reactions, tetrapyrrole metabolism and the Benson-Calvin cycle (Fig. S6A). Additionally, we found a large number of DEGs related to other processes, including hormone metabolism, response to abiotic stress, cell cycle and protein degradation. Among these, we noticed decreased levels of *ATG8A* and *ATG8I*, which encode components of the autophagy conjugation pathway (Fig. 4B and Table S2). Also, multiple transcripts related to auxin metabolism were altered. Particularly, *PIN5* (encoding an auxin transporter that controls auxin levels in the lateral root cap) was decreased, while other transcripts encoding the auxin carriers *PIN1, PIN3*, and *PIN7* were increased; intriguingly, the *LAX3* transcript was also increased (Fig. 4B and Table S2).

Considering the lack of starch observed in *amiR23*-silenced plants treated with NaCl, we focused our attention on DEGs related to starch and sucrose metabolism. The transcripts *PGSIP1* and *PGSIP4*, encoding glycogenin-like proteins putatively involved in the initiation of the starch granule, were reduced. Conversely, three transcripts encoding different ADP-glucose pyrophosphorylase subunits (*APS1, APL3* and *APL4*) and the *GBSS1* transcript (encoding a granule-bound starch synthase) were increased (Fig. 4B and Table S2). We also found that several transcripts encoding enzymes involved in starch degradation were increased, such as *GWD* (or *SEX1*, encoding a glucan, water dikinase) and *SEX4* (encoding a glucan phosphatase), while *BAM9* (encoding a beta-amylase) was decreased (Fig. 4B and Table S2). Transcripts related to sucrose degradation showed different patterns. For instance, *CINV1* and *BFRUCT3*, encoding cytosolic and vacuolar invertases, respectively, were reduced. Conversely, the transcripts encoding the mitochondrial alkaline/neutral invertase C and the cell wall invertase 5 (*CWINV5*) were increased, as well as those encoding sucrose synthases 1 and 3 (*SUS1* and *SUS3*, respectively; Fig. 4B and Table S2).

We then compared the results obtained with *amiR23*-silenced and WT plants treated with NaCl (Fig. S6B). In this case, the changes were less prominent than in the previous analysis (i.e. *amiR23* silenced plants treated with NaCl versus control conditions). These results indicated that some responses to NaCl occurring in WT plants were conserved in *amiR23* silenced plants and were consistent with qPCR data presented in Figs. 3E and 3F. In general, we found DEGs involved in multiple cellular processes, including response to abiotic stress, development, hormone metabolism and protein degradation (Fig. 4B). Some of these transcripts were already observed in the previous comparison. For example, reduced levels of transcripts encoding auxin transporters (*PIN5* and *PIN6*) and increased levels of *ATG8E* (a component of the autophagy pathway). Interestingly, transcripts involved in sucrose and starch metabolism showed the same trend: *CINV1, BFRUCT3* and *BAM9* were reduced, while that encoding the mitochondrial alkaline/neutral invertase C was increased (Fig. 4B and Table S2).

### *amir23*-silenced plants cannot recover from NaCl treatment either with auxin or sucrose

Considering that auxin indirectly promotes the formation of starch granules (Zhang *et al*., 2019), we wondered whether NaCl seriously affecting *amiR23*-silenced plants can be balanced by the effect of auxin. To test this hypothesis, *amiR23*, AT23 and WT plants were grown in normal conditions for five days and then placed in NaCl for 8 h. As previously shown, this treatment reduced starch content in the three genotypes (Fig. 3). The addition of 1 µM IAA (instead of 150 mM NaCl) to the MS medium during 8 h did not affect the scenario observed in control conditions (Fig. 5C), whereas the addition of both 1uM IAA and 150 mM NaCl failed to counter the NaCl effect (Fig. 5D). When the treatment with 150 mM was prolonged for 72 h, both WT and AT23 plants recovered their starch, while *amiR23* plants remained affected (Fig. 5E). The treatment with 1 µM IAA during 72 h did not change the scenario of untreated plants (Fig. 5F), and when both compounds were added together, control and AT23 plants showed starch whereas *amiR23* plants irreversibly lost it (Fig. 5G). The same differences between genotypes were observed when roots were placed in 150 mM NaCl for 8 h and then placed in 1 µM IAA or 1% sucrose for additional 72 h (Figs. 5H and 5H). These results indicated that *amiR23* plants are incapable of recovering from the salinity treatment; therefore AtHB23 plays a crucial in the response to this stress.

**Figure 5.**
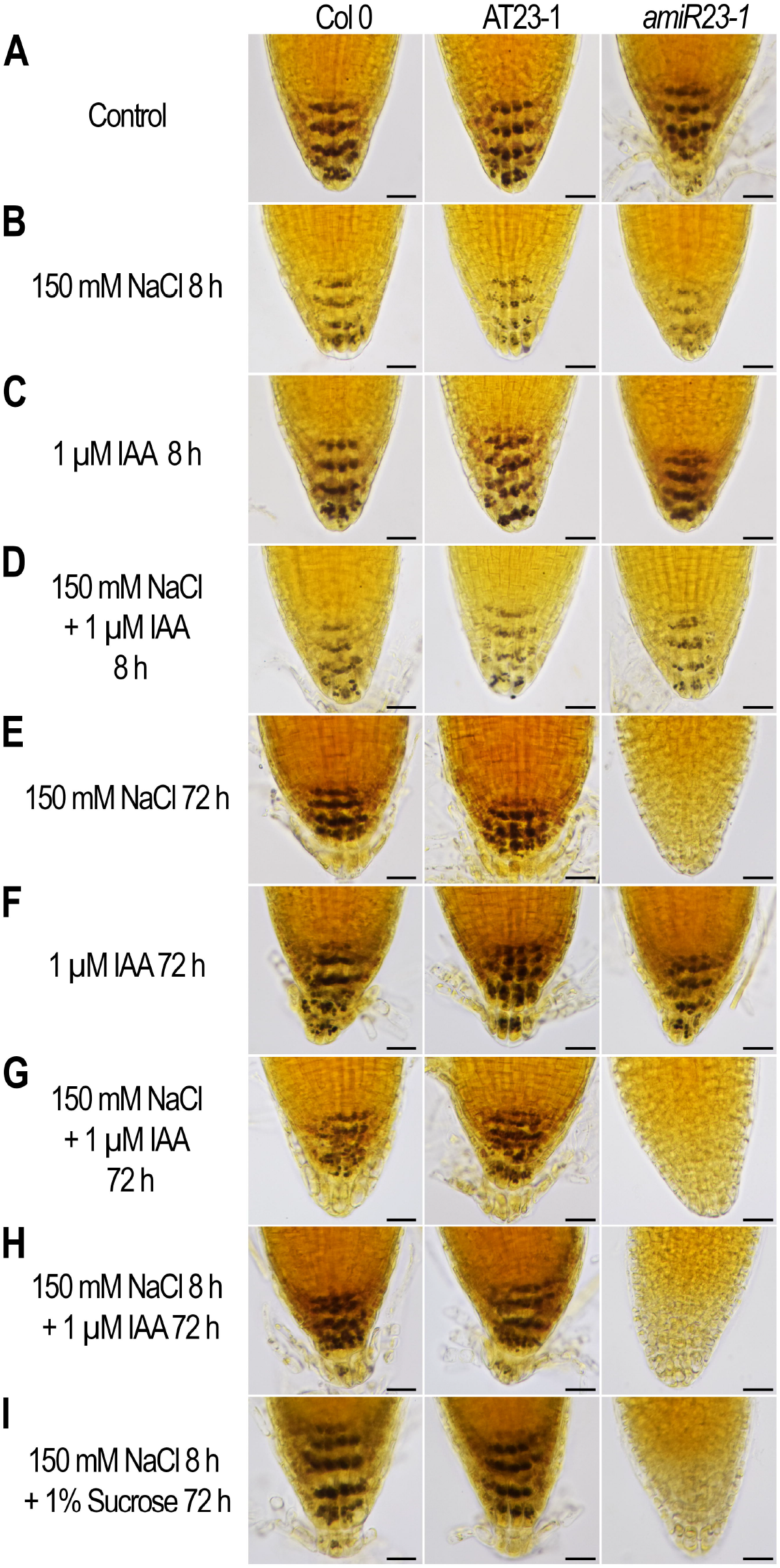
Neither auxin nor sucrose were able to revert amyloplast degradation in *amiR23-silenced* plants. From top to bottom, illustrative pictures of root tips (5-day-old) stained with Lugol solution of Col 0, AT23, and *amiR23* seedlings grown in normal conditions (**A**), after 8 h treatment with 150 mM NaCl (**B**), or with 1 µM IAA(**C**), or with 150 mM NaCl + 1 µM IAA (**D**). (**E**)5-day-old roots treated for 72 h with 150 mM NaCl, or with 1 µM IAA (**F**), or with 150 mM NaCl + 1 µM IAA (**G**), or treated during 8 h with 150 mM NaCl and then transferred to MS medium supplemented with 1 µM IAA (**H**) or 1% sucrose (**I**) for additional 72 h. The assays were repeated at least three times with N: 15/genotype. Black bars represent 50 µm.

### The AtHB23-target gene *LAX3* modulates the main root development in response to NaCl

Considering that *LAX3* and *LBD16* genes were reported as direct targets of AtHB23 in the context of LR development (Perotti *et al*., 2019), we wondered whether these genes were involved in the response to salinity. Transgenic plants bearing the promoters of *LAX3* and *LBD16* fused to the *GUS* reporter gene were analyzed in control conditions and after treatments with 75 mM NaCl in the main, secondary, and tertiary roots. That is because the modulation of these genes was found to be dependent on the root order (Perotti *et al*., 2020). *LAX3* was expressed in the root tip of the main, lateral, and third order roots in 14-day-old seedlings (Fig. 6A), whereas *LBD16* was expressed in the tip of LR and third order root but not in that of the main root even after NaCl treatment (Fig. S7). Surprisingly, after NaCl treatment, the signal corresponding to *LAX3* only disappeared from the tip of the main root, remaining without changes in secondary and tertiary root tips (Fig. 6A). Moreover, transcript levels of *LAX3* measured after the treatment showed a marked reduction, in agreement with histological observations (Fig. 6B). Thus, we compared by histological observation 10-day-old *prLAX3:GUS* and crossed *prLAX3:GUS* × *amiR23-1* seedlings. Unexpectedly, *LAX3* signal disappeared from the main root tip of crossed plants, either treated or untreated (Fig. 6C), hinting at an AtHB23-mediated activation of *LAX3* transcription in the main root. These results strongly suggested that the repression exerted by NaCl on *LAX3* in the main root tip is stronger than the induction promoted by AtHB23, at least in this specific developmental stage and condition.

**Figure 6.**
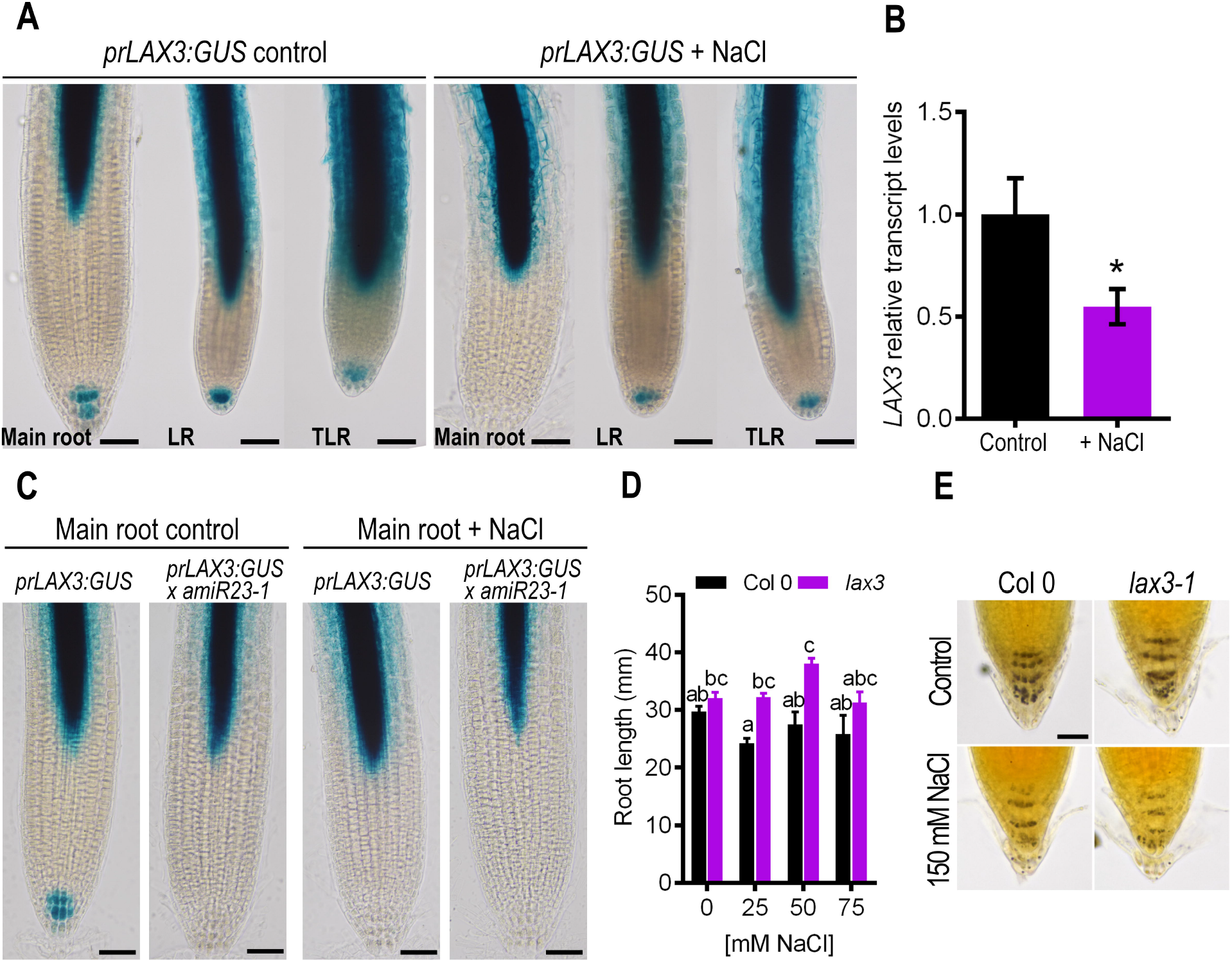
The expression of the auxin carrier *LAX3* is repressed by AtHB23 in front of salt stress in the main root but not in higher order roots. (**A**) Illustrative pictures showing GUS expression driven by *LAX3* promoter *(proLAX3:GUS)* in 14-day-old seedlings. Plants were grown in normal conditions for 3 days and then placed in MS plates (left panel) or MS supplemented with 75 mM NaCl (right panel) for 11 additional days. LR: lateral roots; TLR: tertiary roots. (**B**) Transcript levels of *LAX3* in 5-day-old seedlings grown in normal conditions (control) or treated with 150 mM NaCl for 8 h. The values were normalized with that obtained in normal conditions. Bars represent SEM. (**C**) GUS staining in primary roots of *prLAX3:GUS* or *prLAX3:GUS × amiR23-1* 10-day-old plants grown in normal conditions (left panel) or sowed in MS during 3 days and treated with 75 mM NaCl during 7 additional days (right panel). Black bars represent 100 µm. (**D**) Relative main root length assessed in *lax3* mutants compared with the Col 0 controls, both treated with different NaCl concentrations. Bars represent SEM. Data were analyzed using a two-way ANOVA considering genotype and treatment. Different letters indicate significant differences (Tukey test, P < 0.01). (**E**) Illustrative pictures of Col 0 and *lax3* mutants root tips stained with Lugol in normal conditions or treated with 150 mM NaCl for 8 h.

Then, we wondered whether *LAX3* could alter the root phenotype in NaCl conditions. To this end, *lax3* mutant plants were evaluated after treatments with different NaCl concentrations, ranging from 0 to 75 mM. In control conditions, *lax3* mutants had slightly longer main roots than WT plants, and this difference increased in 25 and 50 mM NaCl (analyzed for 10 days; Fig. 6D). On the other hand, the amount of starch in *lax3* mutant seedlings did not show significant differences with the WT plants, either under control conditions or after the salinity treatment (Fig. 6E). Altogether, these observations indicated that exposed to salinity, LAX3 does not play a crucial role and is probably modulated by different TFs.

## Discussion

Root developmental plasticity determines the capacity of plants to adapt to different soil types. Among many characteristics defining soil quality, salinity emerges as a strong feature affecting plant growth. Moreover, salinity depends on water availability, continuously challenging roots to display their skills to capture water and micronutrients.

Root architecture plasticity involves the development of the main, lateral, and roots of higher order. The development of the main and lateral roots often exhibit opposite responses involving different, sometimes contrary, growth patterns. For example, when subjected to water deficit, the main root grows deeper in order to reach water, and LRs arrest their development (Julkowska and Testerink, 2015).

Under control conditions, root development is governed antagonistically by auxin and cytokinin, but cell division and LR morphogenesis rely, in different ways, on auxin signaling (Petricka *et al*., 2012). For example, in the *diageotropica/wol* mutants (tomato and Arabidopsis) LR development is arrested, and auxin cannot rescue the phenotype (Parizot *et al*., 2008; Ivanchenko *et al*., 2006). It has been reported that mild salinity enhances the growth of the main and LRs, while NaCl increasing levels are detrimental for both, indicating a salt concentration dependency of root growth (Zolla *et al*., 2010; Julkowska *et al*., 2014a). However, in the Col 0 accession, the development of the main root is more affected by salinity than that of LR and depends on the exposure time (Julkowska *et al*., 2014b). Moreover, it seems that the best strategy for roots to survive salinity is to have few but longer LRs to exclude Na^+^ (Julkowska and Testerink, 2015).

In this work, through the study of the homeodomain-leucine zipper transcription factor AtHB23, we exemplify the significant changes that take place in the Arabidopsis genetic program to deal with salinity stress. Previously, it was shown that in normal growth conditions, the knock-down of *AtHB23* provoked an enhancement in LR initiation. This TF is upregulated by ARF7/19, induced the expression of *LAX3* and repressed that of *LBD16* in LR, and, importantly, this genetic program differed between second and third-order roots (Perotti *et al*., 2019, 2020). Under salinity stress, plants with altered levels of *AtHB23* exhibited significant changes in root architecture. The LR phenotype of *amiR23* silenced and AT23 plants was in accordance with the regulation by NaCl (Figs. 1 and 2). Surprisingly, the main root of *AtHB23* knocked-down plants was more affected than LR although we did not detect *AtHB23* expression. Remarkably, the main root arrest, observed in *amiR23* plants, correlates with a compromised survival under high salinity (Fig. 2). The analysis of plants transformed with a shorter segment of the promoter region fused to *GUS* revealed a strong expression in columella cells and an induction by NaCl both in the main and LR, explaining the differential phenotype (Fig. 1). In a salinity environment, *AtHB23*-silenced plants showed arrested growth of the main root and an increment in LRP, stronger than in normal media. The opposite phenotype was exhibited in *AtHB23* overexpressor plants. Interestingly, in the model legume *Medicago truncatula*, the HD-Zip I TF MtHB1 was shown to block LR emergence in response to high salinity by directly repressing the expression of the auxin-responsive LBD TF-encoding gene *MtLBD1*. Moreover, the main root tip of *mthb1* mutants experienced a severe disorganization under stress (Ariel *et al*., 2010a and b). Although the closest homologs of MtHB1 in Arabidopsis are AtHB7 and 12, our results indicate that AtHB23 and MtHB1 may exert similar roles in these distant species.

One of the characteristics observed was the waiving of *amiR23* roots, indicating a loss of the gravitropic response. Gravitropism is a crucial process allowing roots to grow downwards and potentially capturing more water. This phenomenon involves the accumulation of starch granules as statoliths that perceive gravity. This important route is mediated by auxin and affected by salinity conditions (Zhang *et al*., 2019). Although some steps of this process were already described (Iglesias *et al*., 2010), the relationship between salinity, auxin, starch biosynthesis and degradation, and gravity perception remains largely unknown. In normal growth conditions, PIN directional auxin transporters generate an auxin gradient within the root apex, altering the expression of *SS4, PGM*, and *ADG1*, genes encoding enzymes involved in the synthesis of starch granules (Zhang *et al*., 2019).

Degradation of amyloplasts in the columella cells was observed in water-stressed roots, provoked by sorbitol or mannitol (Takahashi *et al*., 2003). The silencing of *AtHB23* resulted in the irreversible loss of starch in the columella cells of the main root in response to NaCl, indicating a crucial role for this TF in the salinity response (Fig. 3). The RNA-Seq data for *amiR23*-silenced plants treated with NaCl showed reduced levels of transcripts encoding different isoforms of extensins and components of the cellulose synthase complex, whereas transcripts encoding several expansins were increased (Figs. 4 and S6). These results suggest altered turnover of the cell wall (reduced synthesis and increased degradation), which could affect its integrity, thus increasing the susceptibility to NaCl (Liu *et al*., 2021). These observations were in accordance with the low survival rate exhibited by *amiR23*-silenced plants after NaCl treatment (Fig. 2).

In agreement with the observed phenotype, starch turnover was severely affected by AtHB23. This was observed not only by individual evaluation of transcripts by RT-qPCR, but also by the transcriptome analysis. Transcripts encoding starch-synthesizing enzymes were decreased, while those encoding starch-degrading enzymes were increased (Figs. 3 and S5). Intriguingly, *ADG1* and *GBSS1* were increased in the RNA-Seq when comparing *amiR23*-silenced plants in control versus salinity conditions (Figs. 4 and S6). These results seem counterintuitive; however, it must be considered that *ADG1* and *GBSS1* were induced in *amiR23*-silenced plants treated with NaCl, but to a lesser extent than in WT and AT23 plants (Figs. 3F and S5). Consequently, these transcripts were not found in the RNA-Seq as DEGS in salinity conditions (*amiR23* versus WT; Figs. 4 and S6). We also noticed reduced levels of the transcripts *PGSIP1* and *PGSIP4* in *amiR23* plants treated with NaCl compared with control conditions (Figs. 4 and 6). Interestingly, Arabidopsis plants expressing a *PGSIP1* RNAi construct showed reduced levels of starch in leaves (Chatterjee *et al*., 2005), which could explain the starchless phenotype of *amiR23* plants treated with NaCl.

Sucrose degradation is catalyzed by sucrose synthases or invertases (Barratt *et al*., 2009). Arabidopsis has six genes encoding cytosolic (*SUS1-4*) and vascular (*SUS5-6*) sucrose synthases and multiple genes coding for cytosolic, vacuolar, and apoplastic invertases. The *sus1/sus2/sus3/sus4* quadruple mutant showed normal contents of sucrose and starch, while the *sus5/sus6* double mutant exhibited reduced callose. Notably, when the genes encoding the main cytosolic invertase isoforms in roots (*CINV1/CINV2*) were mutated, plants exhibited a severely affected phenotype, including loss of starch in the columella cells (Barratt *et al*., 2009; Pignocchi *et al*., 2021). Although the altered morphology of *cinv1/cinv2* mutants was similar to that observed in *amiR23* silenced plants treated with NaCl, transcript levels of *CINV1* and *CINV2* did not significantly change in *amiR23* plants exposed to 150 mM NaCl for 8 h, suggesting that *AtHB23* is not regulating sucrose degradation through this pathway. However, we observed decreased levels of the *CINV1* transcript in *amiR23*-silenced plants exposed to 75 mM NaCl for 6 days (Figs. 4 and S6).

Vacuolar invertases are involved in the regulation of plant growth and development in response to abiotic stress. The tea *INV5* was reported as induced by cold and carbohydrates; the overexpression of this gene promoted taproot and LR elongation involving carbohydrates and auxin signaling (Qian *et al*., 2018). Interestingly, we found consistently reduced levels of the *BFRUCT3* transcript, encoding the vacuolar invertase 1 (Sergeeva *et al*., 2006), in *amiR23* plants (Figs. 4 and S6).

Notably, and considering that auxin promotes starch granule formation, the hormone did not change the amount of starch in columella cells when roots were treated alone neither rescued starch degradation in *amiR23* plants, even after 72 h. Such plants irreversibly lost their starch granules (Fig. 5). These results are in good agreement with those reported by Pignocchi *et al*. (2021), who showed that disruption of auxin signaling does not explain the phenotype of the *cinv1/cinv2* double mutant (see above).

Auxin homeostasis emerged also as an additional factor linked to AtHB23-mediated response to salinity. *LAX3*, previously shown to be directly regulated by AtHB23 (Perotti *et al*., 2019), was repressed by 75 mM NaCl in the main root tip, but not in the tips of higher-order roots (Fig. 6A). While the *LAX3* signal disappeared in the root tip of *prLAX3:GUS × amiR23-1* crosses, indicating a positive regulation by AtHB23, unexpectedly, NaCl repressed *LAX3* expression (Fig. 6C). This puzzling scenario cannot be easily explained. We can hypothesize that although AtHB23 induces *LAX3*, and the former is upregulated by 75 mM NaCl, the impact of salinity overpowers that of this TF. A possible pathway for such effect could be the upregulation of another/unknown HD-Zip I TF, able to heterodimerize with AtHB23, avoiding its action on *LAX3*. In this sense, HD-Zip I TFs are able to heterodimerize (Capella *et al*., 2014), and several members are expressed in the columella cells (Perotti *et al*., 2021). An alternative explanation could be supported on NaCl concentration and treatment time-dependency of the response. *LAX3* expression was repressed in 14-day old crossed plants and in 5-day-old plants treated for 8 h with 150 mM NaCl (Fig. 6C), whereas in the transcriptome analysis, performed with 9-day-old plants subjected for 6 days to a lower salt concentration (75 mM), this gene appeared slightly induced (Table S2). Altogether, the results shown here indicated that AtHB23 plays opposite roles in the main and LR when plants were subjected to salinity and that it is necessary for plant survival under stress (Fig. 7). This TF is required to recover starch biosynthesis after NaCl treatment in order to form granules, necessary to sense and respond to gravity. AtHB23 integrates the modulation of genes participating both in starch turnover and auxin transport.

**Figure 7.**
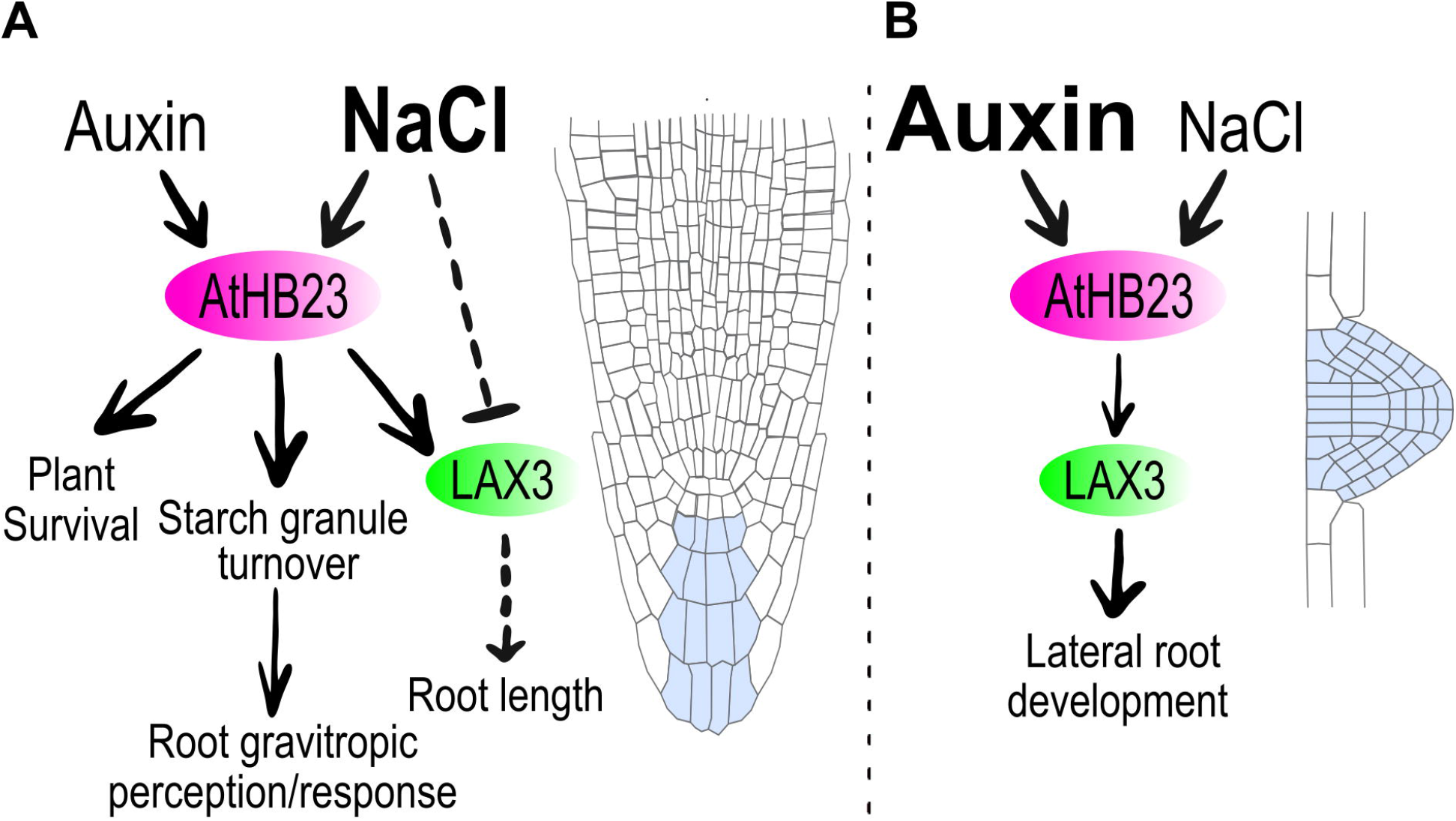
Proposed model for AtHB23 in salinity conditions. Summary of the observations done, focused in the regulation by NaCl and auxin of main (**A**) and lateral (**B**) root development by AtHB23.

## Abbreviations

HD-Zip: homeodomain-leucine zipper
TF: transcription factor
COL 0: wild type
GUS: β-glucuronidase

## Acknowledgements

This work was supported by Agencia Nacional de Promoción Científica y Tecnológica (PICT 2017 0305 and PICT 2019 01916 to RLC) and CONICET. MFP is a CONICET Ph.D. Fellow. ALA, FDA, CMF, and RLC are CONICET Career members.

The authors thank Dr Ranjan Swarup for kindly providing *LAX3* promoter fused to GUS seeds used in this study to our colleague Dr. Javier Moreno.

The authors also acknowledge Dr. Carolina Leimgruber, Dr. Virginia Juárez (UNC), and Dr. Cesar González (IBB-UNER) for technical assistance in TEM analyses.

## Authors’ contribution

Conceived the experiments: MFP, CMF, FDA, and RLC. MFP performed most of the experiments. ALA performed the bioinformatics analyses and wrote the corresponding items. Conceived and wrote the paper: RLC. All authors revised, discussed, and approved the manuscript.

## Data availability

All the data and materials that support the findings of this study are available upon request from the corresponding author.

## Supporting information

Additional Supporting Information may be found in the online version of this article.

**Table S1: oligonucleotides used in this work**

**Table S2: Transcriptomic analysis of wild type and *amiR23* plants in control and salinity conditions**

**Fig. S1. The expression pattern of *AtHB23* is modulated by NaCl in lateral roots**

**Fig. S2. *AtHB23* expression is induced by salinity conditions in the main and lateral roots**

**Fig. S3. *AtHB23* levels affect main root phenotypes**

**Fig. S4. Starch loss in independent *amiR23*-silenced lines under salinity stress is rescued in *amir23x*AT23 plants**

**Fig. S5. Transcript levels of genes involved in starch metabolism turnover**

**Fig. S6. Mapman analysis of the DEGs *amiR23* plants compared with Col 0 in normal or salinity conditions**

**Fig. S7. *LBD16* expression is not altered by salinity conditions**

## Legends to figures

**Figure 1. The expression pattern of *AtHB23* is differentially affected by NaCl in the main and lateral roots, altering whole root development**

(**A-D**) Relative primary root length (**A**) of 8-day-old Col 0 and *amiR23-1* silenced plants, relative density of lateral root primordium (LRP, **B**), lateral root (LR, **c**), and total LR (LRP + LR, **D**) calculated as the number of LRP or LR/mm of main primary root, grown in control conditions during 8 days (control) or 3 days and treated with NaCl (50 or 75 mM), during 5 additional days. (**E-H**) The same as in (**A-D**) evaluated in Col 0 or AT23 (*AtHB23* overexpressor) plants. The values were normalized with those measured in the Col 0 control, taken as 1 (one). All the assays were repeated at least three times with N: 15/genotype. (I) Illustrative pictures showing GUS expression driven by *AtHB23* promoter (short version, *prAtHB23*_*S*_*:GUS*) in the tips of main and lateral roots of 8-day-old seedlings grown in control conditions. (**J**) The same as in (**I**) in plants grown 3 days in normal conditions and then placed during 5 additional days in 75 mM NaCl. LR are emerged lateral roots. Black bar represents 50 µm.

**Figure 2. AtHB23 promoted main root elongation under salinity stress**

(**A**) Main root length evaluated in Col 0, AT23-1 (overexpressors), and *amiR23-1* (silenced) seedlings grown in normal conditions. In the left panel quantitative measurements performed from day 4 after sowing and until day 10, and in the right panel, illustrative pictures of the same roots taken the 10^th^ day. The same plants as in (**A**) were transferred 3 days after sowing to plates supplemented with 25 mM (**B**), 50 mM (**C**) or 75 mM (**D**) NaCl. The assays were repeated at least three times with N: 15/genotype. Black bar represents 1 cm. Different letters indicate significant differences (Tukey test, P < 0.01). Error bars represent SEM.

(**E-F**) Survival rate of plants (Col 0, AT23 and *amiR23*) placed in plates with 100 mM NaCl, 3 days after sawing during the 6 (**E**), and 15 (**F**) additional days. Yellow columns indicate the % of dead plants whereas green ones the % of survivors. The assays were repeated at least three times with N: 15/genotype.

**Figure 3. The amyloplasts in the root tip of *amiR23* silenced plants were degraded in NaCl medium and were unable to recover**

(**A**) From top to bottom, illustrative pictures of root tips (5-day-old) stained with Lugol solution of Col 0, AT23, and *amiR23* seedlings grown in normal conditions (top), after 8 (upper-middle) or 72 h (bottom) treatment with 150 mM NaCl, and then transferred to normal conditions for additional 72 h (lower-middle). Black bar represents 50 µm. (**B**) Illustrative pictures taken with Transmission electron microscopy (TEM) of root tips of 8-day-old seedlings grown in normal conditions (top), treated during 8 h with NaCl (middle), and placed again in normal conditions for 72 h additional hours (bottom). White bars represent 2 µm. (**C**) Starch quantification by ImageJ software in the roots treated or not with 150 mM NaCl. (**D**) Schematic representation of the metabolic pathways for starch synthesis or degradation. (**E-F**) Transcript levels of key genes in Col 0, AT23-1, and *amiR23-1* plants grown in normal conditions during 5 days, treated 8 h with 150 mM NaCl and placed to recover in MS medium during additional 72 h. Genes assessed participating from degradation were *GWD* and *BAM1* (**E**), and from synthesis *PGM*, and *ADG1*. All the values were normalized with the one obtained in Col 0. Bars represent SEM. Data were analyzed using a two-way ANOVA considering genotype and treatment. Different letters indicate significant differences (Tukey test, P < 0.01). The accession numbers of the tested genes are the following: AT5G48300 (ADG1), AT5G51820 (PGM), AT1G10760 (GWD), AT3G23920 (BAM1).

**Figure 4. Transcriptome analyses indicated a differential regulation in *amiR23* roots of genes involved and starch metabolism under salinity**

(**A**) Venn Diagram showing the overlap of the sets of differentially expressed genes (FDR-adjusted p-value < 0.05) in the four different contrasts analyzed, in which the level of one factor is fixed and the two levels of the other factor are compared, e.g. the genotype is Col 0 and the comparison is performed between salinity and control conditions. (**B**) Heatmap showing the expression pattern of relevant genes (rows) in relation to the *amiR23* phenotype. They were selected based on the results of the Mapman analysis of DEGs (Fig. S6), and the inspection of the complete transcriptome (Table S2). The functional annotation of genes is highlighted on the left. Only the three contrasts with a relevant number of DEGs were included (columns).

**Figure 5. Neither auxin nor sucrose were able to revert amyloplast degradation in *amiR23*-silenced plants**

From top to bottom, illustrative pictures of root tips (5-day-old) stained with Lugol solution of Col 0, AT23, and *amiR23* seedlings grown in normal conditions (**A**), after 8 h treatment with 150 mM NaCl (**B**), or with 1 µM IAA (**C**), or with 150 mM NaCl + 1 µM IAA (**D**). (**E**) 5-day-old roots treated for 72 h with 150 mM NaCl, or with 1 µM IAA (**F**), or with 150 mM NaCl + 1 µM IAA (**G**), or treated during 8 h with 150 mM NaCl and then transferred to MS medium supplemented with 1 µM IAA (**H**) or 1% sucrose (**i**) for additional 72 h. The assays were repeated at least three times with N: 15/genotype. Black bars represent 50 µm.

**Figure 6. The expression of the auxin carrier *LAX3* is repressed by AtHB23 under salinity stress in the main root but not in higher order roots**

(**A**) Illustrative pictures showing GUS expression driven by *LAX3* promoter (*proLAX3:GUS*) in 14-day-old seedlings. Plants were grown in normal conditions for 3 days and then placed in MS plates (left panel) or MS supplemented with 75 mM NaCl (right panel) for 11 additional days. LR: lateral roots; TLR: tertiary roots. (**B**) Transcript levels of *LAX3* in 5-day-old seedlings grown in normal conditions (control) or treated with 150 mM NaCl for 8 h. The values were normalized with that obtained in normal conditions. Bars represent SEM. (**C**) GUS staining in primary roots of *prLAX3:GUS* or *prLAX3:GUS* × *amiR23* 10-day-old plants grown in normal conditions (left panel) or sowed in MS during 3 days and treated with 75 mM NaCl during 7 additional days (right panel). Black bars represent 100 µm. (**D**) Relative main root length assessed in *lax3* mutants compared with the Col 0 controls, both treated with different NaCl concentrations. Bars represent SEM. Data were analyzed using a two-way ANOVA considering genotype and treatment. Different letters indicate significant differences (Tukey test, P < 0.01). (**E**) Illustrative pictures of Col 0 and *lax3* mutants root tips stained with Lugol in normal conditions or treated with 150 mM NaCl for 8 h.

**Figure 7. Proposed model for AtHB23 in salinity conditions**

Summary of the observations done, focused in the regulation by NaCl and auxin of main (**A**) and lateral (**B**) root development by AtHB23.

